# Immune differentiation regulator p100 tunes NF-κB responses to TNF

**DOI:** 10.1101/414425

**Authors:** Budhaditya Chatterjee, Payel Roy, Uday Aditya Sarkar, Yashika Ratra, Meenakshi Chawla, James Gomes, Soumen Basak

## Abstract

Stringent regulation of TNF signaling prevents aberrant inflammation. TNF engages the canonical NF-κB pathway for activating the RelA:p50 heterodimer, which mediates specific expressions of pro-inflammatory and immune response genes. Importantly, the NF-κB system discriminates between time-varied TNF inputs. Negative feedback regulators of the canonical pathway, including IκBα, are thought to ensure transient RelA:p50 responses to brief TNF stimulations. The noncanonical NF-κB pathway controls a separate RelB activity associated with immune differentiation. In a systems modeling approach, we uncovered an unexpected role of p100, a constituent of the noncanonical pathway, in TNF signaling. Brief TNF stimulation of p100-deficient cells produced an additional late NF-κB activity consisted of the RelB:p50 heterodimer, which distorted the TNF-induced gene-expression program. Periodic TNF pulses augmented this RelB:p50 activity, which was reinforced by NF-κB-dependent RelB synthesis. In sum, the NF-κB system seems to engage distantly related molecules for enforcing dynamical and gene controls of immune-activating TNF signaling.

## Introduction

Tumor necrosis factor (TNF) is a pleiotropic cytokine whose primary physiological function involves coordinating innate and adaptive immune responses (Kalliolias and Ivashkiv, 2016). TNF activates the canonical NF-κB pathway (Mitchell et al., 2016). The RelA:p50 NF-κB heterodimer is sequestered in the cytoplasm of the unstimulated cells by the inhibitor of κB (IκB) α, β and ε proteins. In the canonical pathway, the IκB kinase (IKK) complex consisting of NEMO and IKK2 (or IKKβ) phosphorylates IκBs that promotes their degradation and nuclear activation of RelA:p50. Once in the nucleus, RelA:p50 mediates the expression of pro-inflammatory and immune response genes.

Typically, TNF briefly stimulates tissue resident cells due to its short half-life *in vivo* (Beutler et al., 1985). Previous studies demonstrated that the NF-κB system, in fact, distinguishes between brief and chronic TNF signals. Brief TNF stimulation induces a transient RelA:p50 activity peak persisting in the nucleus for about an hour (Hoffmann et al., 2002; Werner et al., 2008). For varied TNF concentration, chronic TNF stimulation triggers an additional second wave of protracted RelA:p50 activity, which displays oscillatory behaviour at single-cell resolution (Cheong et al., 2006; Hoffmann et al., 2002; Nelson et al., 2004). Importantly, RelA:p50 also induces the synthesis of the inhibitors of the canonical pathway, including IκBα, IκBε and A20 (Lee et al., 2000; Scott et al., 1993). A series of elegant studies (Ashall et al., 2009; Kearns et al., 2006; Paszek et al., 2010; Werner et al., 2008) suggested that coordinated functioning of these negative feedback regulators determines dynamical RelA:p50 responses to time-varied TNF inputs. Indeed, it is thought that RelA:p50 regulation by the canonical NF-κB pathway largely provides for dynamically controlled expressions of specific pro-inflammatory and immune response genes during TNF signaling (Hoffmann, 2016). On the other hand, deregulated TNF signaling has been implicated in several human ailments, including inflammatory bowel disorders and neoplastic diseases (Kalliolias and Ivashkiv, 2016).

The noncanonical NF-κB pathway controls a separate RelB NF-κB activity. In resting cells, p100 encoded by *Nfkb2* retains RelB in the cytoplasm (Sun, 2012). Noncanonical signaling induced by B-cell activating factor (BAFF) or lymphotoxin α_1_β_2_ (LTα_1_β_2_) activates the NF-κB inducing kinase (NIK), which in association with IKK1 (or IKKα) phosphorylates p100. Subsequently, proteasome removes the C-terminal inhibitory domain from p100 that generates the mature p52 NF-κB subunit and releases the RelB:p52 NF-κB heterodimer into the nucleus. The noncanonical pathway elicits a weak but sustained RelB activity, which selectively drives the expression of genes involved in the differentiation of immune cells and the development of immune organs. Notably, p100 deficiency was shown to produce a minor RelB:p50 containing nuclear NF-κB activity, which partially compensated for the absence of immune-organogenic RelB:p52 functions in *Nfkb2*^*-/-*^ mice (Basak et al., 2008; Derudder et al., 2003; Lo et al., 2006).

Although TNF *per se* does not activate noncanonical signaling, molecular connectivities between the canonical and noncanonical NF-κB pathways exist (Mitchell et al., 2016). For example, TNF-activated canonical signaling induces the expression of genes encoding RelB and p100 from the respective NF-κB target promoters (Sun, 2012). Albeit weakly, p100 binds to RelA (Basak et al., 2007; Tao et al., 2014) and IκBα interacts with RelB (Roy et al., 2017; Shih et al., 2012). More so, RelA and RelB heterodimers were shown to possess overlapping as well as distinct DNA binding and gene-expression specificities in specialized cells (de Oliveira et al., 2016; Roy et al., 2017; Shih et al., 2012; Siggers et al., 2011; Zhao et al., 2014). We asked if these interconnectivities between canonical and noncanonical NF-κB modules contribute to TNF signaling.

Here, we identified that the absence of the noncanonical signal transducer p100 obliterated the capacity of the NF-κB system to discriminate between brief and chronic TNF signals. Brief TNF treatment, akin to chronic simulations, induced a biphasic NF-κB response in p100-deficient cells. However, the late NF-κB DNA binding activity induced in p100-deficient cells was consisted of the RelB:p50 heterodimer, which distorted TNF-mediated gene control. Mechanistically, NF-κB-driven RelB synthesis reinforced the RelB:p50 activity in TNF-treated p100-null cells. In sum, our study suggests that an interlinked NF-κB system, and not solely the components of the canonical module, engages distantly related molecular species with seemingly distinct biological functions to ensure specific transcriptional responses to time-varied TNF inputs.

## Results

### A mathematical model of the integrated NF-κB system predicts a role of p100 in TNF signaling

Mathematical reconstructions of cellular networks offer insights on the underlying signal-processing mechanisms (Basak et al., 2012; Le Novere, 2015). The NF-κB system consists of interlinked canonical and noncanonical modules (Fig 1a). To understand how the integrated NF-κB system preserves dose-duration control of TNF signaling, we utilized previously published mathematical model (Roy et al., 2017), which depicted interconnected canonical and noncanonical NF-κB pathways, subsequent to necessary revisions (see Supplementary Information for details). Theoretical IKK2 activity profiles of varying peak amplitude or duration were used as model inputs (Fig 1b, Fig S1a and Fig S1b). Our computational simulations broadly captured the previously described NF-κB signaling dynamics (Hoffmann et al., 2002; Werner et al., 2008). For example, the duration of nuclear NF-κB (NF-κBn) response was insensitive to changes in the amplitude of IKK2 signal, but proportionately increased as a function of the duration of IKK2 input (Fig 1c). Consistent to previous reports (Hoffmann et al., 2002), our computational model revealed a role of IκBα in this dynamical control (Fig 1d). Interestingly, our computational analyses revealed an aberrant dynamical control of NF-κBn in the *Nfkb2*-deficient system with IKK2 activities of varied duration producing comparable ∼6-8 hr of responses (Fig 1d).

**Figure 1:**
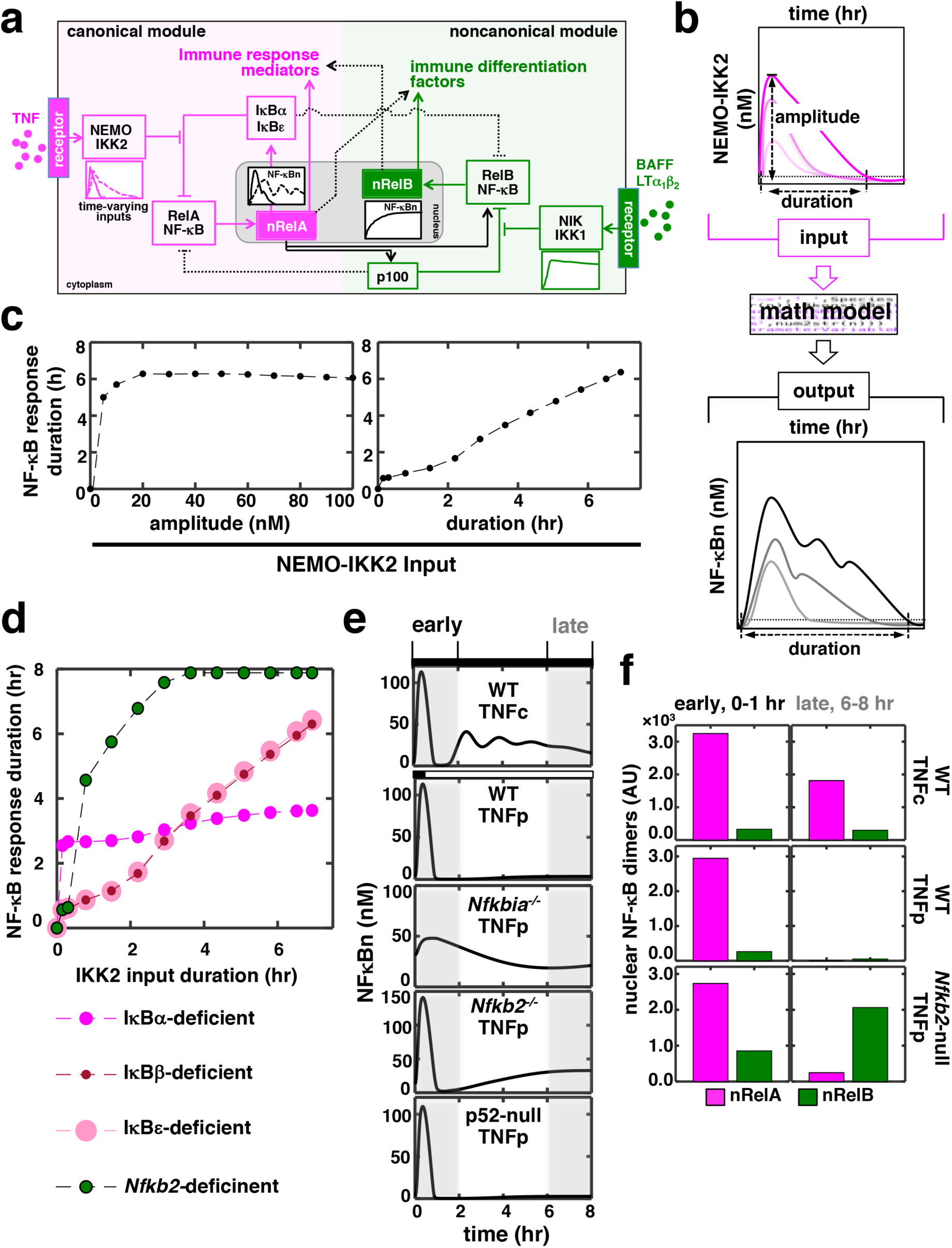
*In silico* studies identify a role of p100 in discriminating between time-varying TNF inputs. **a)** A graphical depiction of the NF-κB system. TNF through the canonical pathway (magenta) dynamically regulates the activity of the RelA:p50 heterodimer, which mediates the expression of immune response genes. BAFF or LTα_1_β _2_ induces a distinct RelB NF-κB activity via a separate noncanonical pathway (green) for driving the expression of immune differentiation factors. However, these two NF-κB pathways are molecularly connected and display certain overlap in relation to gene expressions. Solid and dotted black lines represent major cross-regulatory mechanisms and those involving less-preferred biochemical reactions, respectively. NF-κBn, nuclear NF-κB activity. nRelA and nRelB represent corresponding nuclear heterodimers. **b)** Schema describing *in silico* production function analyses. Briefly, theoretical IKK2 activity profiles of various peak amplitudes and durations were fed into the mathematical model, and NF-κBn responses were simulated in a time-course. Durations were estimated as the time elapsed above a specific threshold value, which was determined as the sum of the basal NF-κB or IKK activity and 5% of the corresponding basal-corrected peak activity, in the corresponding activity curves. **c)** and **d)** Graph plot of the duration of simulated NF-κBn responses as a function of the peak amplitude or the duration of theoretical IKK2 inputs. IKK2 activities of various peak amplitude but with invariant 8 hr of duration (c, left) or with various durations but identical 60nM peak amplitude (c, right and d) were used. Computational simulations involved **c)** the WT system and **d)** the indicated mutant systems. **e)** *In silico* studies revealing NF-κBn responses in a time-course in WT and various mutant systems. Experimentally derived IKK2 activity profiles, obtained using MEFs treated with TNF either chronically (TNFc) or for 0.5 hr (TNFp), were used as model inputs. Early (0-2 hr) and late (6-8 hr) phases have been marked using grey boxes. **f**) Computational modeling predicting TNFp-induced nuclear activities of RelA and RelB heterodimers in WT and *Nfkb2*-null systems. Early and late activities were determined as the area under the corresponding activity curve between 0-2 hr and 6-8 hr, respectively, subsequent to correction for basal values. AU, arbitrary unit.

In experimental settings, chronic (TNFc) TNF treatment of mouse embryonic fibroblasts (MEFs) generates a long-lasting IKK2 activity persisting for ∼ 8 hr, while brief0.5 hr (TNF pulse, TNFp) of TNF stimulation elicits only ∼1 hr of IKK2 signal (Shih et al., 2009; Werner et al., 2008). When we fed these experimental IKK2 activity profiles into our mathematical model as inputs (Fig S1c), our computational simulations efficiently recapitulated experimental NF-κBn responses (Hoffmann et al., 2002; Shih et al., 2009; Werner et al., 2008). For example, the long-lasting IKK2 activity associated with TNFc triggered a prolonged, biphasic NF-κBn response composed of the RelA:p50 heterodimer (Fig 1e and 1f). Short-lived IKK2 input related to TNFp produced only a transient 1hr of NF-κBn response in the WT system. As expected, a weakened negative feedback extended the TNFp-induced NF-κB response beyond 1hr in the IκBα-deficient system. Corroborating our computational studies involving theoretical IKK2 inputs, simulation of the TNFp regime in the *Nfkb2*-deficient system indeed produced a prolonged NF-κBn response, whose temporal profile was somewhat comparable to that of the TNFc-induced NF-κBn activity. The prolonged activity induced in the *Nfkb2*-deficent system was biphasic where the late phase lasted for ∼8 hr (Fig 1e). However, this late activity was absent in the p52-null system, where conversion of p100 into p52 was not permitted. Finally, our simulation studies indicated that signal-induced nuclear accumulation of the alternate RelB:p50 heterodimer generated this late-acting NF-κBn response to TNFp in the *Nfkb2*-deficient system (Fig 1f). Therefore, our mathematical modeling studies predicted a role of the non-canonical signal transducer p100 in producing appropriate NF-κB responses to time-varying TNF inputs.

### p100 restrains late-acting RelB:p50 NF-κB response to brief TNF stimulation

To verify experimentally the predictions of our mathematical model, we treated MEFs with TNF and measured the resultant NF-κBn activities in a time-course using the electrophoretic mobility shift assay (EMSA). TNFc treatment of WT cells induced a biphasic NF-κBn response comprising of an early peak, which lasted for ∼1 hr, and a gradually weakening second phase persisting between 3-8 hr (Fig 2a). TNFp treatment of WT MEFs produced 1hr of the early peak activity, which was substantially broadened in TNFp-treated *Nfkbia*^*-/-*^ MEFs lacking IκBα (Fig 2b). As predicted by our computational model, TNFp induced a prolonged NF-κBn response in *Nfkb2*^*-/-*^ MEFs that consisted of the early peak and a progressively strengthening second phase (Fig 2b). Of note, TNFc induced a similar biphasic activity in *Nfkb2*^*-/-*^ cells (Fig S2a). Our shift-ablation assay confirmed that the TNFp-induced, late NF-κBn DNA binding activity was composed of the RelA:p50 heterodimer in *Nfkbia*^*-/-*^ cells. As described (Basak et al., 2008), we noticed in *Nfkb2*^*-/-*^ MEFs a low level of basal RelB:p50 activity, which was reinforced upon TNFp treatment and produced the robust, late-acting NF-κB response (Fig 2b and 2c). This late RelB:p50 activity persisted in the nucleus of *Nfkb2*^*-/-*^ MEFs even 16 hr after TNFp stimulation (Fig S2b). TNFp treatment of 10 min or 20 min duration, which produce ∼30 min of IKK2 activity (Werner et al., 2008), efficiently induced the late NF-κBn response in the absence of p100 (Fig 2d). Finally, IL-1β, which induces NF-κB signaling transiently in WT cells (Werner et al., 2008), produced a similar late RelB:p50 activity in *Nfkb2*^*-/-*^ MEFs (Fig 2e and Fig S2c). Our studies suggested that p100 enforced dynamical NF-κB control by preventing the late-acting RelB:p50 response to short-lived IKK2 signals generated by pro-inflammatory cytokines. However, deficiency of p100 and that of the well-articulated negative feedback regulator, IκBα caused distinct aberrations with respect to the temporal profile and the composition of the signal-induced nuclear NF-κB activity.

**Figure 2:**
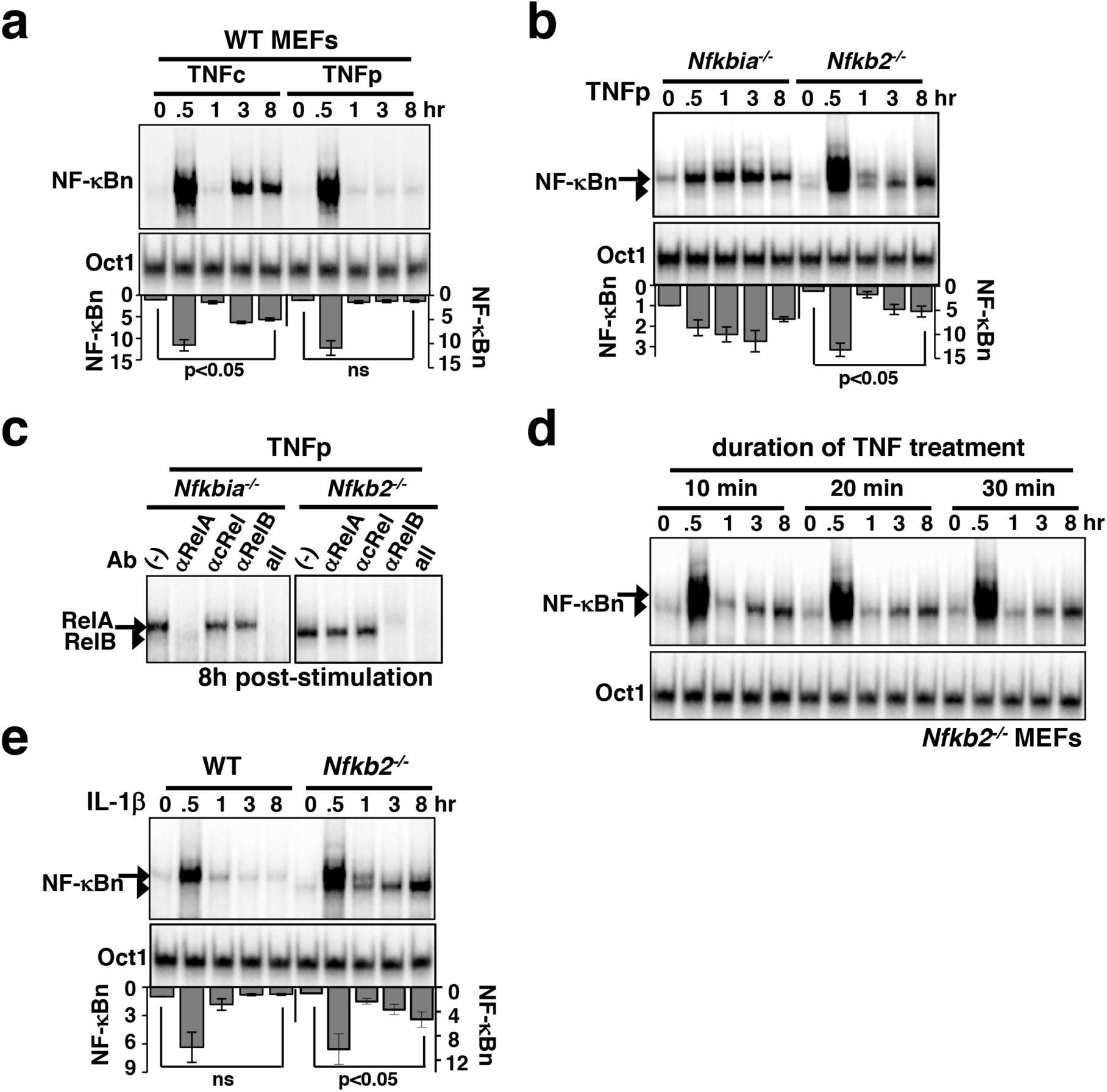
Brief TNF stimulation of *Nfkb2*^*-/-*^ MEFs induce an additional late NF-κB activity composed of the RelB:p50 heterodimer. **a)** WT MEFs were subjected to TNFc or TNFp treatments, cells were harvested at the indicated time-points after the commencement of stimulations, and NF-κBn DNA binding activities were resolved in EMSA (top panel). DNA binding activity of Oct1 served as a loading control (middle panel). Bottom: signals corresponding to NF-κBn were quantified from four independent experiments and presented in relation to the basal activity in a bargraph. Statistical significance was evaluated using Student’s *t* test. **b)** EMSA comparing NF-κBn induced in a time-course in *Nfkbia*^*-/-*^ and *Nfkb2*^*-/-*^ MEFs upon TNFp treatment (top panel). As determined in Fig 2C, the arrow and the arrowhead indicate RelA-containing and RelB-containing NF-κB complexes, respectively. Bottom: quantitative analysis of the total NF-κBn activities from four experimental replicates. **c)** Composition of the NF-κBn activities that persisted after 8 hr of TNFp treatment in *Nfkbia*^*-/-*^ and *Nfkb2*^*-/-*^ MEFs, was determined in the shift-ablation assay. Antibodies against the indicated NF-κB subunits were used for ablating the respective DNA binding complexes in EMSA. Data represents two independent experiments. **d)** EMSA demonstrating NF-κBn induced in a time-course in *Nfkb2*^*-/-*^ MEFs subjected to brief TNF stimulation for 10 min, 20 min or 30 min (top panel). Data represents two biological replicates. **e)**Time-course analysis of NF-κB DNA binding activity induced upon IL-1β treatment of WT or *Nfkb2*^*-/-*^ MEFs (top panel). Bottom: quantified NF-κB signal intensities; data represent four experimental replicates. Quantified data presented in this figure are means ± SEM.

### Dissecting molecular mechanisms underlying the late RelB:p50 response to brief TNF stimulation in the absence of p100

Utilizing a variance-based, multiparametric sensitivity analysis method (Chatterjee et al., 2016), we investigated the biochemical mechanism underlying late-acting RelB:p50 response to TNFp in the *Nfkb2*-deficient system. We assembled the large number of model parameters into 48 distinct groups (Fig 3a). Each of these groups consisted of functionally related biochemical parameters associated with a specific molecular species. For instance, kinetic rate parameters associated with the synthesis of IκBα, including constitutive and NF-κB-responsive transcriptions as well as translation, were grouped together. Using Monte Carlo sampling, we explored the parameter space surrounding the initial values simultaneously among the different parameter groups. The effect of parameter uncertainty for individual parameter groups on the late RelB:p50 activity was summarized as the total effect index (Chatterjee et al., 2016). Our analyses revealed that group-V parameters, which included rate parameters associated with NF-κB-driven and constitutive syntheses of *Relb* mRNA as well as translation of *Relb* mRNA into RelB protein, played a dominant role in determining the late RelB:p50 response (Fig 3b and Fig S3a). We distinguished between the parameters belonging to group-V using local sensitivity analysis, which indicated that particularly NF-κB-mediated transcription of *Relb* promoted the late RelB:p50 response to TNFp in the *Nfkb2*-deficient system (Fig 3c). Both RelA and RelB heterodimers are capable of inducing the expression of *Relb* mRNA from the endogenous NF-κB target promoter (Basak et al., 2008; Roy et al., 2017). Our computational studies suggested that RelA-mediated transcription of *Relb* mRNA was not sufficient and RelB-dependent autoregulatory synthesis was required for this late RelB:p50 response in p100-deficient cells (Fig 3d).

**Figure 3:**
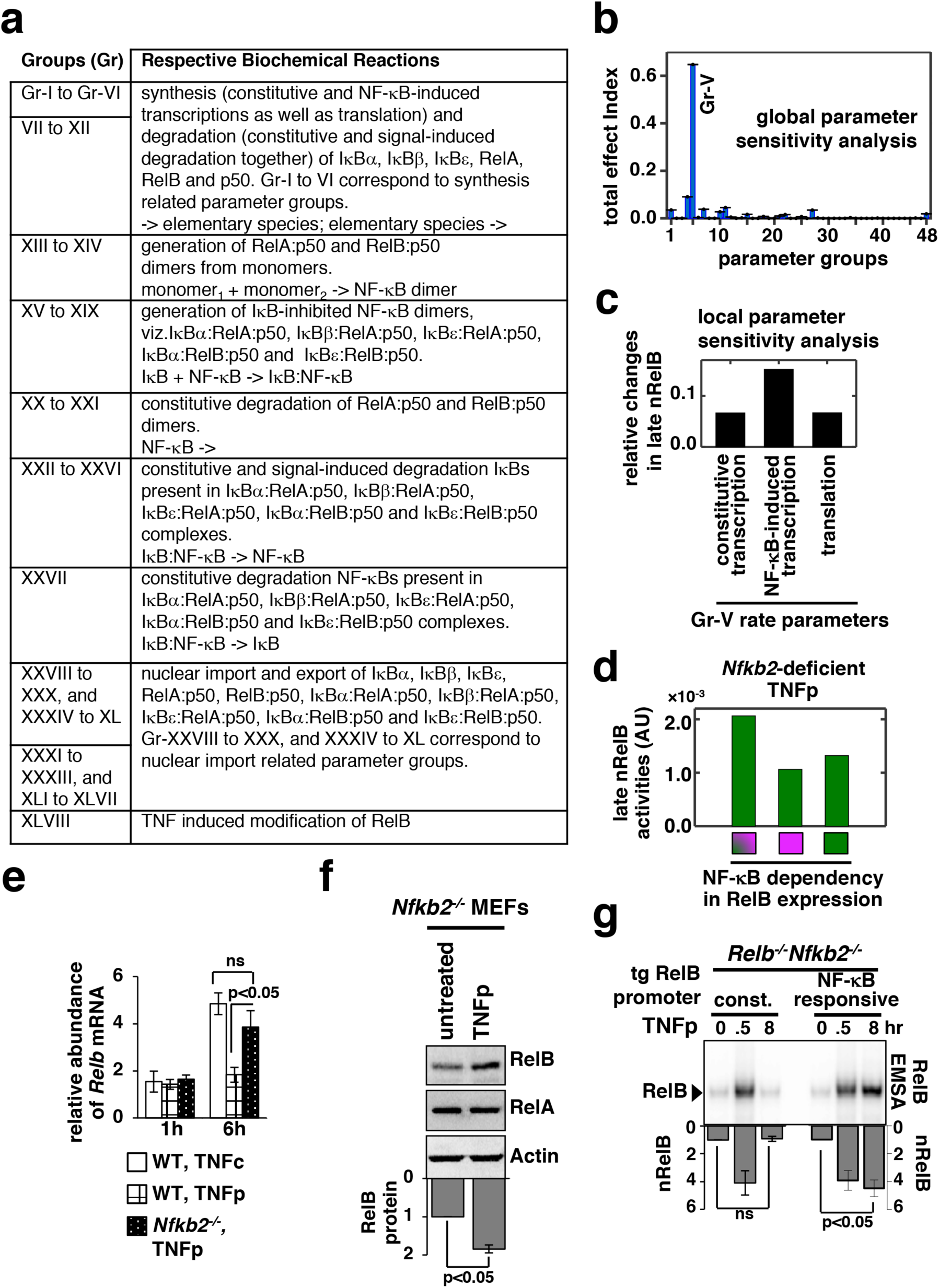
NF-κB-dependent RelB synthesis promotes the late RelB:p50 activity induced upon brief TNF treatment of *Nfkb2*^*-/-*^ cells. **a)** Parameter groups analysed in the variance-based, multiparametric sensitivity analysis. **b)** Variance-based multiparametric analysis revealing the total effect index, which represent the effect of the parameter uncertainty on the late (6-8 hr) RelB:p50 activity induced by TNFp in the *Nfkb2*-deficient system for the individual parameter groups. Standard bootstrapping was used for estimating error ranges. Gr, group. Gr-V consists of parameters related to RelB synthesis; including constitutive and NF-κB induced transcriptions as well as translation. **c)** Local sensitivity analysis revealing the effect of 10% increase in the indicated parameters belonging to Gr-V on the late RelB:p50 response to TNFp in the *Nfkb2*-deficient system. Differences in the basal-corrected, RelB:p50 activity between the unperturbed system and perturbed systems were scored. **d)** Computational simulations of the late nRelB response to TNFp in the *Nfkb2*-deficient system, where the expression of RelB is induced either by RelA as well as RelB heterodimers (indicated with a square with magenta and green) or exclusively by RelA (magenta) or RelB (green). **e)** WT and *Nfkb2*^*-/-*^ MEFs were treated with TNFp before being subjected to qRT-PCR analysis of *Relb* mRNA abundances normalized to that of *Actb* mRNA. WT MEFs treated with TNFc were used as control. Bargraphs demonstrate the abundances of mRNAs in TNF-treated cells relative to those measured in the untreated cells. Data represent four biological replicates. **f)** *Nfkb2*^*-/-*^ MEFs were treated with TNFp and harvested at 6 hr post-stimulation before being analysed by Western blotting. Actin served as a loading control. Bottom: densitometric analysis of the relative abundance of RelB protein in whole cell extracts; data represent five biological replicates. **g)** TNFp-induced nRelB activity in *Relb*^*-/-*^*Nfkb2*^*-/-*^ MEFs stably expressing RelB from a retroviral transgene (tg) either constitutively (const.) or from an NF-κB responsive promoter. Ablating RelA DNA binding with an anti-RelA antibody, residual nRelB activities were revealed by RelB-EMSA. Data represent four independent experiments. Quantified data presented in this figure are means ± SEM.

We tested these computational predictions experimentally. We observed that TNFc activated delayed expression of *Relb* mRNA, which is encoded by a NF-κB target gene, in WT MEFs (Fig 3e). Consistent with the lack of late NF-κBn activity in WT MEFs subjected to TNFp treatment, TNFp-induced expressions of *Relb* mRNA were less prominent in these cells (Fig 3e). However, TNFp treatment of *Nfkb2*^*-/-*^ MEFs strongly induced the synthesis of *Relb* mRNA and protein at 6 hr post-stimulation that temporally coincided with the late RelB activity observed in these cells (Fig 3e and Fig 3f). Furthermore, expression of RelB from an NF-κB insensitive, constitutive promoter in *Relb*^*-/-*^*Nfkb2*^*-/-*^ MEFs was inadequate for triggering the late RelB:p50 response to TNFp (Fig 3g and Fig S3b). Our combined mathematical and biochemical analyses established that NF-κBn-driven RelB expressions promoted progressive nuclear accumulation of the RelB:p50 heterodimer in the absence of p100 in response to TNFp.

### RelB:p50 modifies the TNF-mediated gene-expression program in *Nfkb2*^*-/-*^ MEFs

Previous studies, though conducted using disparate cell types and stimulation regimes, identified both distinct and overlapping control of genes by various NF-κB dimers (Smale, 2012). We asked if the TNF-activated RelB:p50 heterodimer modulated RelA-dependent expressions of immune response genes. We initially focused on the TNFc regime, which induced both RelA:p50 and RelB:p50 DNA binding activities in *Nfkb2*^*-/-*^ MEFs at 6 hr post-stimulation (Fig 4a). We also utilized *Relb*^*-/-*^*Nfkb2*^*-/-*^ and *Rela*^*-/-*^*Nfkb2*^*-/-*^ MEFs those elicited either RelA:p50 or RelB:p50 activities, respectively (Fig 4a and Roy et. al., 2017). As a control, we used *Rela*^*-/-*^*Relb*^*-/-*^*Rel*^*-/-*^ cells, which lacked all three transcription-competent NF-κB subunits RelA, RelB and cRel. In microarray mRNA analysis, these knockout cells were compared with WT MEFs, which induced RelA:p50 DNA binding activity, for TNFc-induced gene-expressions. We considered genes whose expressions were induced at least 1.3 fold at 6 hr upon TNFc treatment in *Nfkb2*^*-/-*^ MEFs, but not in *Rela*^*-/-*^*Relb*^*-/-*^*Rel*^*-/-*^ cells. Based on their differential expressions in *Relb*^*-/-*^*Nfkb2*^*-/-*^ and *Rela*^*-/-*^*Nfkb2*^*-/-*^ MEFs, we catalogued the resultant 304 NF-κB-dependent genes into six distinct clusters, which were arranged further into four gene-groups (Gr-I to Gr-IV; see Fig 4b, Supplementary Information, Materials and Methods). Gr-I and Gr-II genes were induced in WT and *Nfkb2*^*-/-*^ MEFs as well as in *Relb*^*-/-*^ *Nfkb2*^*-/-*^ cells (Fig 4a and Fig 4b). Gr-II, but not Gr-I, genes were induced in *Rela*^*-/-*^*Nfkb2*^*-/-*^ cells (Fig 4a and Fig 4b). Interestingly, TNFc triggered the expression of additional genes belonging to Gr-III and Gr-IV in *Nfkb2*^*-/-*^ cells that were not induced in WT MEFs. *Rela*^*-/-*^ *Nfkb2*^*-/-*^ MEFs, but not *Relb*^*-/-*^*Nfkb2*^*-/-*^ cells, supported Gr-III gene expressions; Gr-IV genes were not induced in either of these two genotypes. Our genetic analyses involving knockout cells suggested that a subset (Gr-II) of the RelA NF-κB-target genes induced by TNFc in WT cells was activated by RelB:p50. But RelB:p50, either alone (Gr-III) or in collaboration with RelA:p50 (Gr-IV), participated in the expression of additional genes in *Nfkb2*^*-/-*^ MEFs that were not normally induced in WT cells in the presence of p100.

**Figure 4:**
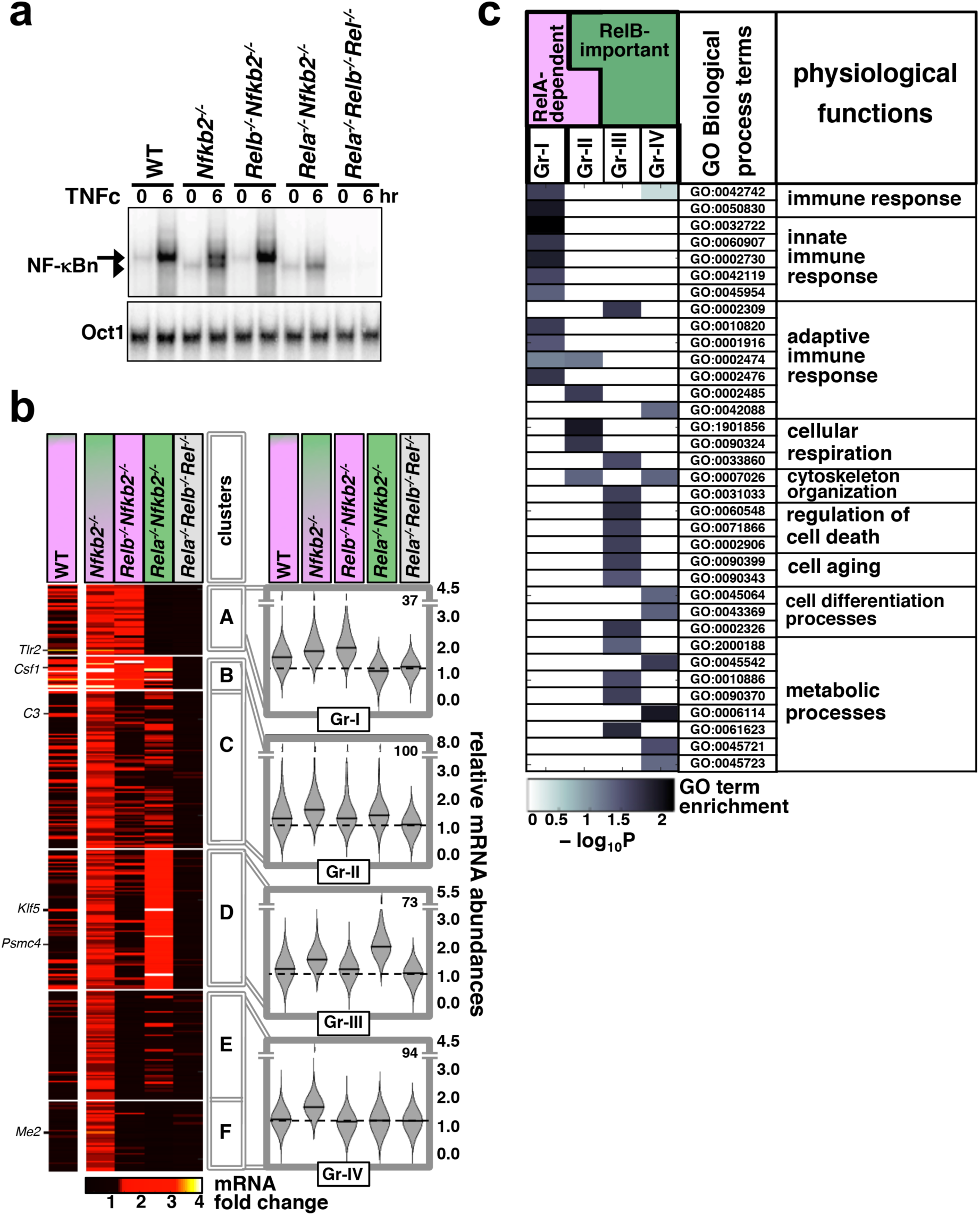
Global analyses reveal distinct gene controls by the TNF-activated RelB:p50 heterodimer. **a)** MEFs of the indicated genotypes were treated with TNFc for 6 hr before being subjected to EMSA. The data represents three independent experiments. **b)** In our microarray mRNA analysis, we considered genes with high confidence detection in biological replicates across various knockout MEFs, and at least 1.3 fold increase in the average expression upon 6 hr of TNFc treatment in *Nfkb2*^*-/-*^ cells, but less than 1.3 fold average induction in *Rela*^*-/-*^*Relb*^*-/-*^*Rel*^*-/-*^ cells to arrive onto a list of 304 genes. Heatmap demonstrates binary logarithmic transform of TNF-induced fold changes in the expressions of these genes in the indicated knockout cells clustered using the partition around medoids algorithm. A representative data using WT MEFs has been indicated in the left column. The resultant six gene-clusters were arranged into four gene-groups. Representative genes belonging to different groups have been indicated. Right: violin plots show relative frequency distributions of fold change values and corresponding medians for various genotypes as well as the number of members in each gene-group. **c)** Functional enrichment of various Gene Ontology for Biological Process terms in the indicated gene-groups was determined by topGO. A subset of highly enriched terms in either of the gene-groups is highlighted. Broad physiological functions associated with these GO terms have been also indicated.

Finally, we subjected these gene-groups to gene ontology (GO) analyses. Consistent with the well-articulated role of the canonical pathway in immune-activating TNF signaling, Gr-I and Gr-II comprising TNFc-induced RelA-dependent genes were enriched for GO terms associated with innate and adaptive immune responses (Fig 4d). Gr-II also scored highly for terms linked to cellular respiration. Gr-III and Gr-IV consisting of RelB-important genes activated in p100-deficient cells were instead enriched for terms associated with cellular differentiation, aging, and cell death as well as metabolic processes. However, these RelB-important genes scored poorly for immune response related GO terms. Taken together, our genome-scale analyses implied that the RelB:p50 heterodimer activated in the absence of p100 modified the TNFc-mediated gene-expression program by inducing the expression of distinct sets of genes encoding functions separate from immune activation.

### p100 determines the specificity and the dynamical control of TNF-mediated gene expressions

It was shown earlier that sustained expressions of NF-κB-dependent genes require prolonged RelA:p50 DNA binding activity, which is induced in WT cells by TNFc (Covert et al., 2005; Hoffmann et al., 2002; Werner et al., 2005). We asked if TNFp stimulation of *Nfkb2*^*-/-*^ MEFs triggered persistent gene expression, particularly those regulated by RelB:p50. To this end, we first validated our genome-scale data by qRT-PCR analyses. Consistently, TNFc treatment for 6 hr triggered the RelA-dependent expression of *Tlr2*, a representative member of Gr-I, in WT and *Nfkb2*^*-/-*^ MEFs (Fig 5a). Either the RelA or the RelB activity was sufficient for the transcription of the NF-κB-target *Csf1* belonging to Gr-II. The RelB activity was necessary for the expression of the Gr-III member *Klf5*, which was induced in *Nfkb2*^*-/-*^ and *Rela*^*-/-*^*Nfkb2*^*-/-*^ MEFs but not WT and *Relb*^*-/-*^*Nfkb2*^*-/-*^ cells. Finally, *Me2*, a Gr-IV member, was induced selectively in *Nfkb2*^*-/-*^ MEFs, which produced both RelA:p50 and RelB:p50 nuclear activities. *Rela*^*-/-*^*Relb*^*-/*^ *Rel*^*-/-*^ cells did not activate these genes.

**Figure 5:**
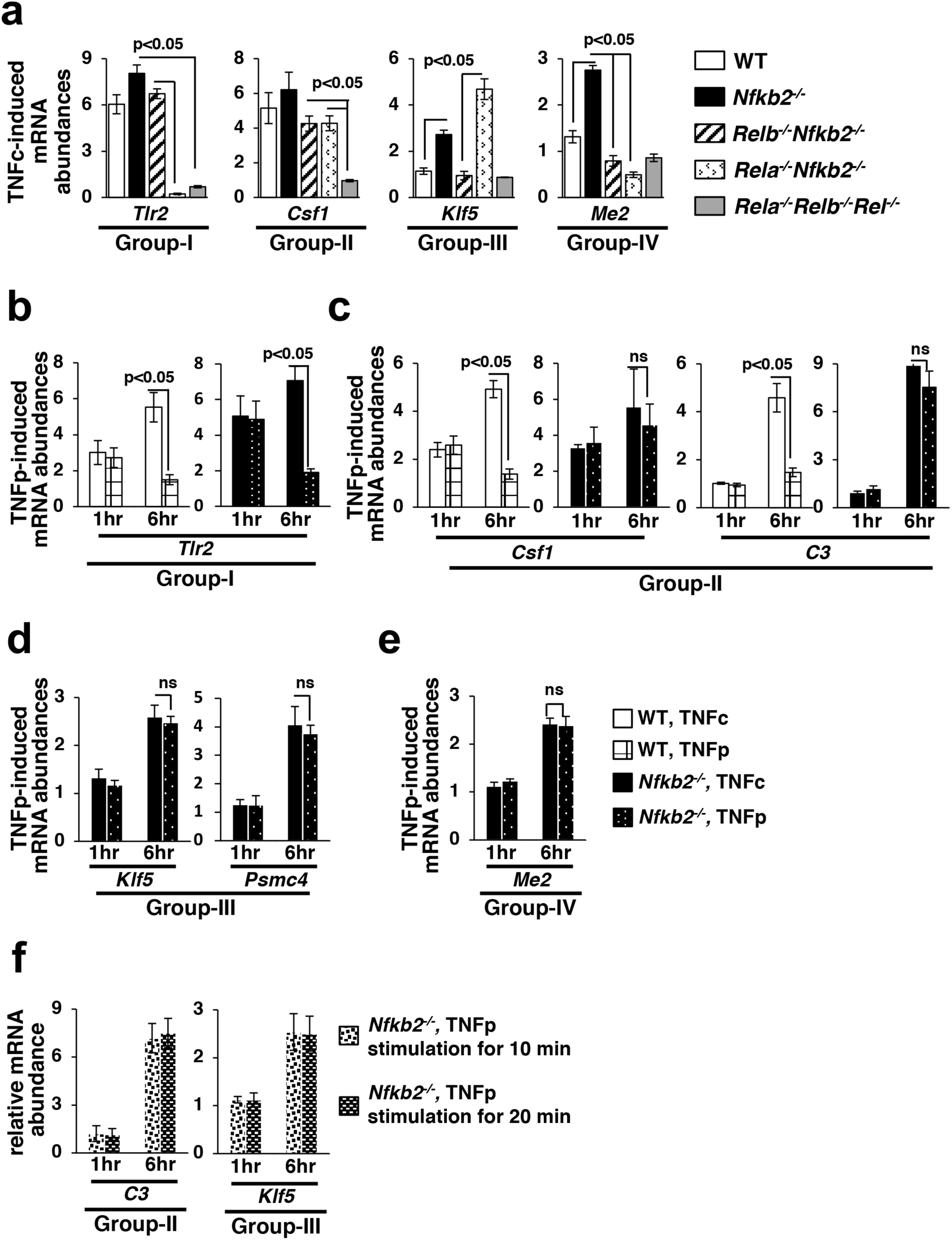
Brief TNF stimulation provokes delayed, RelB-regulated gene expressions in *Nfkb2*^*-/-*^ MEFs. **a)** WT and knockout MEFs were subjected to TNFc treatment for 6h, and the expressions of the indicated genes belonging to different gene-groups were measured by qRT-PCR. Bargraphs demonstrate the abundances of the corresponding mRNAs in stimulated cells relative to those measured in untreated cells. Data represent three biological replicates. **b)** to **e)** WT and *Nfkb2*^*-/-*^ MEFs were subjected to TNFc or briefly stimulated with TNF for 0.5 hr (TNFp), and subsequently cells were harvested at the indicated time-points before being subjected to qRT-PCR analyses. Bargraphs demonstrate TNF-induced expressions of the indicated genes, representing various gene-groups, in relation to untreated cells. Data represent four independent experiments. **f)** *Nfkb2*^*-/-*^ MEFs were treated with TNF briefly for 10 min or 20 min before being subjected to qRT-PCR analyses of the indicated mRNA abundances. Bargraphs demonstrates TNF-induced gene-expressions after 1h and 6h of the commencement of cell stimulation in relation to untreated cells. Data represent four biological replicates. Quantified data presented in this figure are means ± SEM.

We then compared TNFc and TNFp regimes for the expression of genes belonging to various gene-groups. Our time-course analyses demonstrated that TNFc induced progressive accumulation of mRNAs encoding Gr-I and Gr-II genes between 1 hr and 6 hr post-treatment, and that TNFp failed to sustain the expression of these genes at 6 hr in WT cells (Fig 5b and Fig 5c). Indeed, *Nfkb2*^*-/-*^ MEFs upheld the expression of Gr-II genes *Csf1* and *C3* at 6hr post-TNFp stimulation. Moreover, TNFc and TNFp were equally proficient in stimulating the delayed expression of Gr-III genes *Klf5* and *Psmc4* as well as the Gr-IV gene *Me2* in *Nfkb2*^*-/-*^ MEFs (Fig 5d and Fig 5e). Even 10 min of TNF treatment of *Nfkb2*^*-/-*^ cells led to a substantial expression of the RelB-important genes (Fig 5f), which were not activated in WT cells. Collectively, p100 enforced both the dynamical control and the specificity of TNF-induced gene-expressions. Brief TNF stimulation of *Nfkb2*^*-/-*^ MEFs not only sustained the expression of a subset of RelA-dependent genes but also provoked delayed expressions of additional RelB-important genes.

### Repeated pulses of TNF reinforce the late-acting RelB:p50 response in *Nfkb2*^*-/-*^ cells

When administered at short intervals, repeated TNF pulses produce a refractory state in WT cells leading to a deteriorating RelA:p50 peak activity in response to TNF (Adamson et al., 2016; Ashall et al., 2009). We examined the effect of periodic TNF pulses on the late RelB:p50 activity induced in *Nfkb2*^*-/-*^ MEFs. Our computational simulations revealed that a one-hour separation between two consecutive TNF pulses diminished the early RelA:p50 response in both WT and *Nfkb2*-deficient systems, but a separation of 2 hr or more did not show any effect (Fig 6a and Fig 6b). However, the late RelB:p50 activity induced at 8 hr in response to TNFp in the *Nfkb2*-deficient system was further enhanced by a subsequent TNFp for a pulse separation of 1-5 hr. The heightened late RelB activity was accompanied by increased abundances of *Relb* mRNA and protein (Fig S4a). Our experimental analyses substantiated that as compared to a single pulse, two or three consecutive TNF pulses augmented the late RelB activity as well as the abundances of *Relb* mRNA and protein in *Nfkb2*^*-/-*^ MEFs (Fig 6c and Fig S4b). Finally, double or triple TNF pulses enhanced the delayed expression of RelB-important genes in *Nfkb2*^*-/-*^ cells (Fig 6d). These studies identified an important role of p100 in the periodic TNFp stimulation regime; although p100 did not participate in the downregulation of the RelA activity, its absence triggered escalating RelB:p50 NF-κB response.

**Figure 6:**
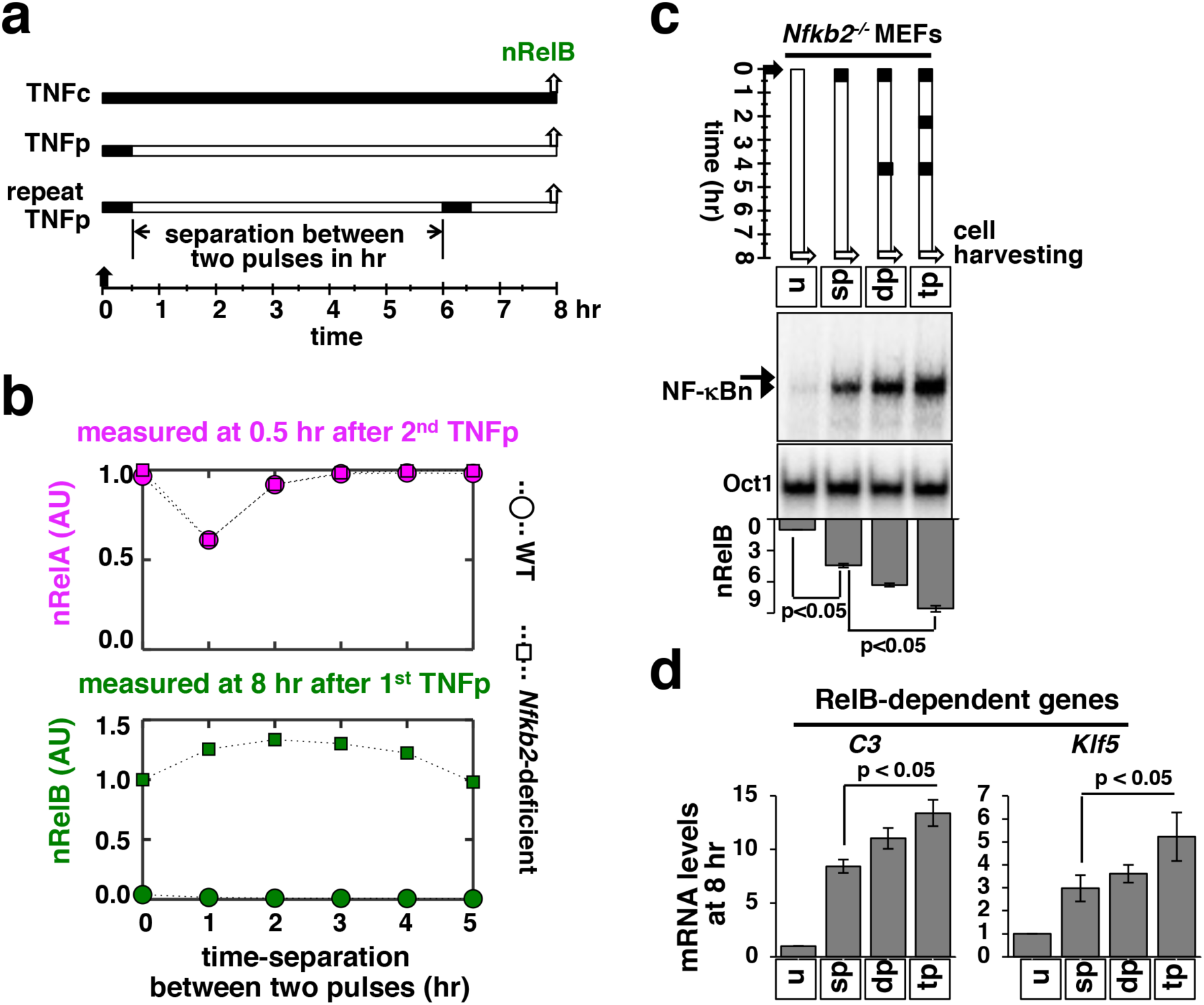
Repeated TNF pulses of *Nfkb2*^*-/-*^ cells strengthen late RelB:p50 signaling. **a)** Schema describing repeated TNFp regime: the short-lived IKK2 input associated with the TNFp regime was fed into the model successively with varied separation time between two TNF pulses and corresponding NF-κBn was simulated. **b)** Computational studies revealing the early nRelA activity induced 0.5 hr after the second TNFp (top) and the late nRelB activity induced 8 hr after the first TNFp (bottom) as a function of the separation time between two successive pulses in WT and *Nfkb2*-deficient systems. The early and late activities were normalized to those induced in response to a single pulse. **c)** *Nfkb2*^*-/-*^ MEFs were treated with either a single TNFp (single pulse, sp) or two successive TNFp separated by 4 hr (double pulse, dp) or three pulses at 2 hr intervals. Cells were harvested 8 hr after the first pulse and analysed for NF-κBn by EMSA. u denotes untreated. Bottom: quantitative analysis of the nRelB activities; data represent three experimental replicates. **d)** *Nfkb2*^*-/-*^ MEFs were subjected to the indicated treatments; cells were harvested 8 hr after the first pulse and expressions of the indicated genes were measured by qRT-PCR. Bargraphs represent three biological replicates. Quantified data presented in this figure are means ± SEM of three biological replicates.

## Discussion

Brief and chronic TNF stimulation elicits transient and prolonged NF-κB activities, respectively, consisting of the RelA:p50 heterodimer, which mediates the expression of specific immune response genes. The NF-κB system distinguishes between time-varied TNF inputs apparently because of the IκBα-mediated negative feedback hardwired in the canonical module (Mitchell et al., 2016). In contrast, p100 transduces the noncanonical NF-κB signal, which mediates nuclear activation of RelB heterodimers during immune differentiation (Sun, 2012). We found an additional function of p100, which discriminates between brief and chronic TNF signals (Fig 7). An absence of p100 provoked a prolonged, biphasic NF-κB response to brief TNF stimulation. However, the late NF-κB activity in *Nfkb2*^*-/-*^ MEFs was composed of RelB:p50, and not RelA:p50 heterodimer. Furthermore, RelB:p50 modified the TNF-induced gene-expression program in *Nfkb2*^*-/-*^ cells. The precursor function of p100 provides for RelB activation during immune differentiation. Our study suggested that p100-mediated inhibition, which insulated RelB from canonical signaling, shaped dynamically controlled specific transcriptional responses to immune-activating TNF.

**Figure 7:**
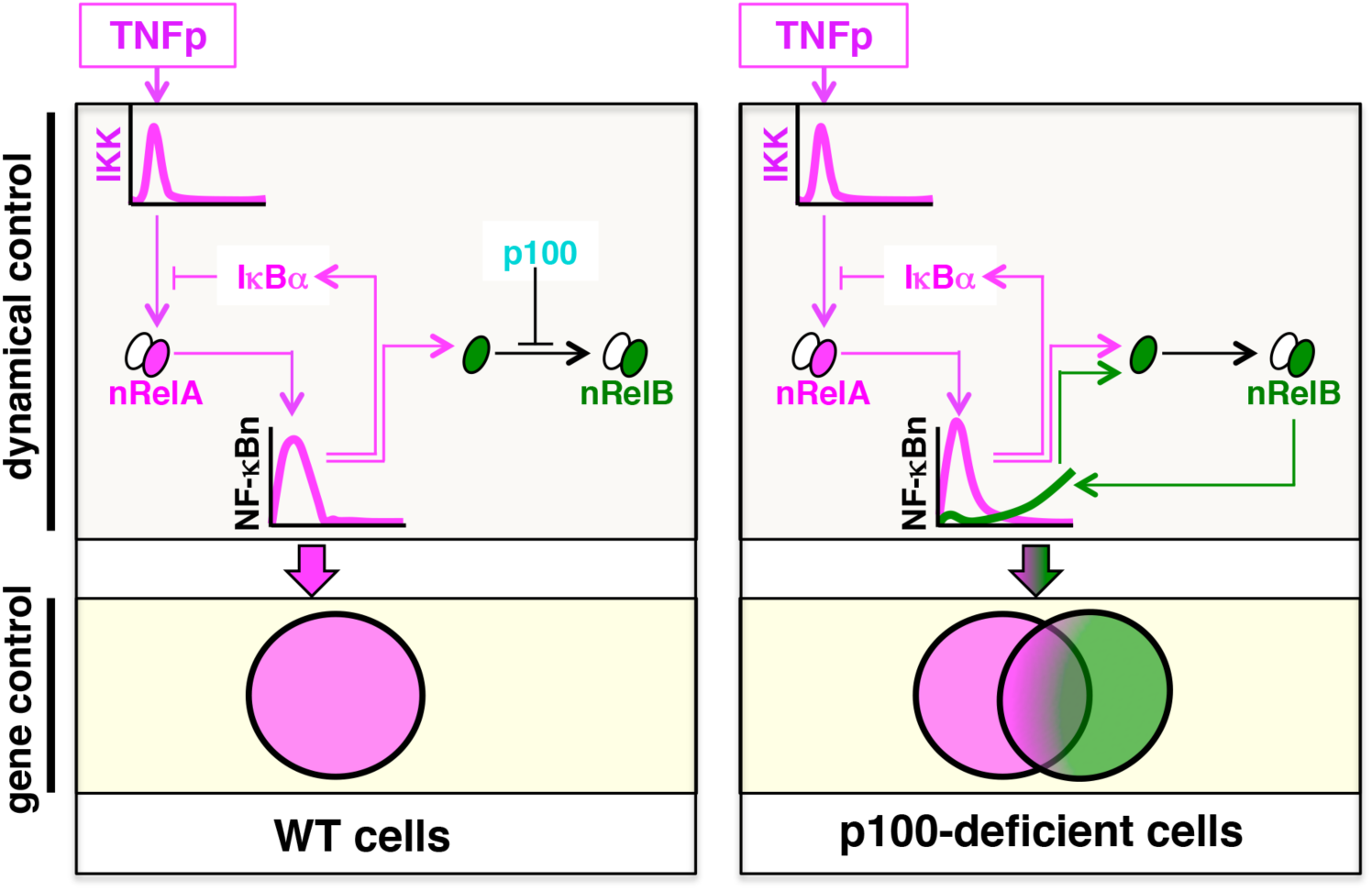
The proposed model explaining the role of p100 in the dynamical control and the gene-expression of specificity of TNF-induced NF-κB signaling. Brief TNF stimulation elicits transient NF-κB activity composed of the RelA:p50 heterodimer, which mediates the expressions of immune response genes. An absence of p100 triggers a late RelB:p50 NF-κB activity in response to TNF that induces also the expressions of genes involved in metabolic and cellular differentiation processes.

As such, TNF and other canonical pathway inducers do not cause degradation of p100, which is proteolyzed during non-canonical signaling. But TNF-activated canonical pathway induces the expression of *Nfkb2* mRNA, and the noncanonical signal transducer p100 also interacts with RelA (Basak et al., 2007; Tao et al., 2014). A previous investigation, which utilized computational model and cells lacking RelB, suggested that p100 is redundant to IκBα in mediating post-induction attenuation of the TNF-induced RelA:p50 NF-κB activity (Shih et al., 2009). Interestingly, it was shown that RelB:p50 binds to IκBα in the absence of p100 and is activated by canonical signaling induced upon chronic TNF treatment (Derudder et al., 2003; Roy et al., 2017; Shih et al., 2012). Our brief TNF stimulation regime instead captured an important role of p100 in preventing late-acting RelB:p50 NF-κB response. Brief TNF stimulation of *Nfkb2*^*-/-*^ cells resulted in a transient RelA:p50 activity, but a prolonged RelB:p50 response. It was shown that an absence of p100 produces a low level of constitutive RelB:p50 activity in *Nfkb2*^*-/-*^ cells (Basak et al., 2008). Our mechanistic analyses clarified that TNF-induced RelB synthesis, including RelA-dependent feedforward and autoregulatory transcriptions, reinforced this RelB:p50 activity eliciting the robust, late-acting NF-κB response in *Nfkb2*^*-/-*^ cells (Fig 7). In other words, restraining synthesis-dependent RelB activation, p100 ensured transient immune-activating RelA NF-κB response to brief TNF stimulation.

RelB-mediated gene control remains partly obscure. RelB was shown to both activate and inhibit the expression of RelA-target genes (Roy et al., 2017; Saccani et al., 2003; Shih et al., 2012; Xia et al., 1997). Our global analyses using genetically tractable MEFs revealed overlapping and non-redundant gene functions of these NF-κB heterodimers. We found that RelB:p50, in fact, supported the expression of a subset of RelA-target immune response genes. Either alone or in collaboration with RelA, RelB induced additional genes in *Nfkb2*^*-/-*^ MEFs. These RelB-important genes were not activated by TNF in WT cells and were related to immune differentiation and metabolic processes. In *Nfkb2*^*-/-*^ MEFs subjected to brief TNF stimulation, RelB:p50 not only sustained the expression of certain RelA-target genes but also induced protracted expressions of these RelB-important genes. Recent *in vitro* DNA interaction studies and ChIP-seq analyses established that the RelA and RelB heterodimers bind to largely similar κB sequences (de Oliveira et al., 2016; Siggers et al., 2011; Zhao et al., 2014). In line with an earlier proposal (Smale, 2012), we speculate that DNA-protein interactions play a rather insignificant role in determining the gene-expression specificity of NF-κB heterodimers, and that the gene-expression specificity is contingent upon the interaction of NF-κB heterodimers with other transcription factors and cofactors. Future ChIP-seq studies ought to elaborate the gene-regulatory mechanism underlying expressions of RelB-important genes in the p100-deficient cell system.

Stimulus-specific cellular responses are often achieved through distinct dynamical control of shared signaling kinases and transcription factors. For example, neuronal growth factor (NGF) induces the sustained activity of extracellular signal-regulated kinase (Erk) for promoting cell differentiation. In contrast, transient Erk activation by epidermal growth factor (EGF) causes cell proliferation (Marshall, 1995; Santos et al., 2007). Genome-wide knockdown studies indicated that a vast regulatory network, and not a handful of components belonging to specific pathways, controls the amplitude of the activity of these signaling molecules (Friedman and Perrimon, 2007). Our study offered evidence that distantly related molecular species within an interconnected cellular network also modulate the duration of signal-responses. In addition to the constituents of the canonical NF-κB pathway, the immune differentiation regulator p100 controlled dynamical inflammatory signaling. A lack of IκBα leads to sustained RelA:p50 NF-κB response upon brief TNF stimulation. Brief TNF stimulation of p100-deficient cells, however, provoked a late RelB:p50 activity, which possessed both overlapping and distinct gene functions in relation to RelA:p50. Therefore, engagement of multiple regulators appears to control the temporal profile and the composition of the NF-κB activity induced by short-lived pro-inflammatory cytokines.

In addition to regulating immune responses, TNF also contributes to immune differentiation (Sedger and McDermott, 2014). Interestingly, the noncanonical signal transducer RelB has been implicated in certain TNF-dependent immune differentiation processes, including osteoclastogenesis (Tanaka and Nakano, 2009; Vaira et al., 2008). We postulate that removal of p100 by immune differentiating cues may provide for a mechanism of cell-type and tissue microenvironment specific tuning of physiological TNF responses involving canonical RelB activity. Despite its detrimental effect on the signal-induced RelA activity, repeated TNF pulses heightened the late-acting RelB:p50 response in p100-deficient cells and augmented the abundance of RelB-important mRNAs encoding metabolic regulators. Interestingly, *Nfkb2* was shown to be frequently mutated in cancers (Courtois and Gilmore, 2006), and both aberrant inflammations, as well as altered metabolism, have been implicated in neoplastic diseases (Dang, 2012). A synthesis-dependent IRF4 activity promoted the survival of multiple myeloma cells (Shaffer et al., 2008). In this context, it will be important to investigate further if the synthesis-driven RelB activity induced in response to TNF pulses provokes abnormal gene-expressions in cancerous cells with dysfunctional p100.

In sum, the NF-κB signaling system is comprised of interlinked canonical and noncanonical modules and controls the activity of multiple transcription factors, which have overlapping as well as distinct gene functions. Our study emphasizes that this interconnected NF-κB system, and not the individual NF-κB signaling modules, directs dynamically controlled activity of the specific NF-κB heterodimers in response to TNF.

## Materials and Methods

### Mice, Cells and Plasmids

WT and gene-deficient C57BL/6 mice were used in accordance with the guidelines of the Institutional Animal Ethics Committee of the National Institute of Immunology (approval no. #258/11). MEFs generated from E13.5 embryos were used either as primary cells or subsequent to immortalization by the 3T3 protocol. *Relb*^*-/-*^*Nfkb2*^*-/-*^ MEFs expressing transgenic RelB from retroviral constructs were reported earlier (Roy et al., 2017).

### Biochemical analyses

In the TNFp regime, MEFs were treated briefly with 1ng/ml of TNF (Roche, Switzerland). Subsequently, TNF-supplemented media was substituted with TNF-free media, and cells were harvested at the indicated times after the commencement of the TNF treatment. In certain instances, cells were subjected to repeated pulses of TNF at the specified time intervals. Alternately, cells were treated chronically with 1ng/ml of TNF (TNFc) or stimulated with 10ng/ml IL-1β (Biosource, USA). As described (Banoth et al., 2015), nuclear and whole cell extracts were analysed by EMSA and Western blotting, respectively. The gel images were acquired using PhosphorImager (GE Amersham, UK) and quantified in ImageQuant 5.2.

### Gene expression analyses

Total RNA was isolated from MEFs, stimulated either briefly or chronically with 10ng/ml of TNF, using RNeasy kit (Qiagen, Germany). A detailed description of microarray mRNA analyses is available in the Supplementary Information. The partition around medoid-based clustering analysis (Reynolds et al., 2006) was implemented in the Cluster package in R; the heatmap and violin plots were generated in MATLAB. For determining the significance of gene-expression differences between various genotypes within a given gene-group, we conducted multiple hypotheses testing and computed the effect sizes (Supplementary Information). See Table S1 for a description of genes belonging to different gene-groups. The MIAME version of the microarray data is available on NCBI-GEO (accession no. GSE68615). The enrichment of the Gene Ontology terms was determined by Fisher’s exact test using the ‘weight algorithm’ available in topGO (Alexa et al., 2006) and the entire Illumina MouseRef-8 v2.0 gene-array was used as the background. qRT-PCR was performed as described earlier (Roy et al., 2017); see Table S2 for the description of primers.

### Computational modeling

We utilized a previously published mass action kinetics-based NF-κB mathematical model (Roy et al., 2017) subsequent to necessary refinements (Supplementary Information). These refinements improved the performance of the model with respect to the *Nfkbia*-deficient system, but preserved the model behaviour observed earlier in WT and *Nfkb2*-deficient systems (Roy et al., 2017). The model was stimulated using Ode15s in MATLAB (2014b, Mathworks, USA). The abundances of various molecular species during early signaling was determined as the area under the respective timecourse curves between 0-2 hr, and those during late signaling was estimated between 6-8 hr. Variance-based, multiparametric sensitivity analysis has been described (Chatterjee et al., 2016). Using iterative Monte Carlo sampling (1000 simulations), we simultaneously explored a predetermined range (±10%) of parameter space around the initial values for the indicated parameter groups. The parameters belonging to a specific group were altered by the same factor for a given simulation.

### Statistical analysis

Error bars were shown as S.E.M. of 3-6 experimental replicates. Quantified data are means ± SEM, and two-tailed Student’s t-test was used for verifying statistical significance unless otherwise mentioned. Statistical tests associated with global analyses have been detailed in the Supplementary Information.

## Supplemental information

Supplemental Information includes four figures and two tables associated with the main text, a detailed description of global gene expression analyses as well as mathematical modelling studies.

## Author contributions

BC carried out *in silico* studies under the supervision of SB and JG. PR conducted cell-based analyses with the assistance from UAS, YR and MC and the guidance from SB. BC and PR wrote the manuscript with SB. The authors declare no conflict of interest.

## Acknowledgements

We thank members of the Systems Immunology Laboratory (SIL) for critical discussions; V. Kumar, SIL, NII for technical help; P. Nagarajan from SAF, NII for help with animal husbandry. Research in the PI’s laboratory is funded by an intermediate fellowship (500094/Z/09/Z) to SB from Wellcome Trust DBT India Alliance, and SERB, Department of Science and Technology, Govt. of India (EMR/2015/000658) and NII-core. PR thanks CSIR, BC thanks UGC, YR and MC thank DBT for research fellowships.

